# Infection of primary nasal epithelial cells differentiates among lethal and seasonal human coronaviruses

**DOI:** 10.1101/2022.10.17.512617

**Authors:** Clayton J. Otter, Alejandra Fausto, Li Hui Tan, Noam A. Cohen, Susan R. Weiss

## Abstract

The nasal epithelium is the initial entry portal and primary barrier to infection by all human coronaviruses (HCoVs). We utilize primary nasal epithelial cells grown at air-liquid interface, which recapitulate the heterogeneous cellular population as well as mucociliary clearance functions of the *in vivo* nasal epithelium, to compare lethal (SARS-CoV-2 and MERS-CoV) and seasonal (HCoV-NL63 and HCoV-229E) HCoVs. All four HCoVs replicate productively in nasal cultures but diverge significantly in terms of cytotoxicity induced following infection, as the seasonal HCoVs as well as SARS-CoV-2 cause cellular cytotoxicity as well as epithelial barrier disruption, while MERS-CoV does not. Treatment of nasal cultures with type 2 cytokine IL-13 to mimic asthmatic airways differentially impacts HCoV replication, enhancing MERS-CoV replication but reducing that of SARS-CoV-2 and HCoV-NL63. This study highlights diversity among HCoVs during infection of the nasal epithelium, which is likely to influence downstream infection outcomes such as disease severity and transmissibility.

## INTRODUCTION

To date, seven human coronaviruses (HCoVs) are known to infect humans, causing a range of respiratory disease (Coleman and Frieman, 2014; Kesheh *et al*., 2022). Three of these HCoVs emerged from animal reservoirs to cause significant public emergencies in the past 20 years and have been categorized as lethal HCoVs due to their propensity to cause life-threatening pneumonia in infected patients (Wang, Grunewald and Perlman, 2020). Severe acute respiratory syndrome (SARS)-CoV first appeared in South China and caused an epidemic beginning in 2002 that resulted in a total of 8,422 infections and 916 deaths (case-fatality rate 11%) (Chan-Yeung *et al*., 2003; Li *et al*., 2020). Middle East respiratory syndrome-CoV (MERS-CoV) was first identified in Saudi Arabia in 2012 and has caused over 2500 cases and 894 deaths (case-fatality rate of 34.5%) (Zaki *et al*., 2012; *MERS-CoV Worldwide Overview*, 2022; *MERS Situation Update*, 2022). Most recently, SARS-CoV-2, the agent responsible for coronavirus disease 2019 (COVID-19) resulted in the ongoing global pandemic that has caused over 570 million cases and 6.3 million deaths (as of 7/27/22) (*World Health Organization COVID19 Dashboard*, 2022). Four additional HCoVs (HCoV-NL63, -229E, -OC43, and HKU1) infect humans, circulate seasonally (causing 15-25% of common cold cases), and are generally associated with less severe respiratory disease (Wat, 2004; Gaunt *et al*., 2010). Importantly, while the seasonal/common HCoVs typically cause self-limiting upper respiratory tract infections in humans, they can cause more severe disease and lower respiratory tract infections in at-risk populations such as neonates, the elderly, and immunocompromised individuals (Chiu *et al*., 2005).

CoVs have been classified into distinct genera based on serology and phylogenetic clustering. All three lethal HCoVs (SARS-CoV, MERS-CoV, and SARS-CoV-2) and two of the nonlethal HCoVs (HCoV-OC43, -HKU1) are betacoronaviruses, while HCoV-229E and HCoV-NL63 are alphacoronaviruses (Zhou, Qiu and Ge, 2021). CoVs of all genera are enveloped, non-segmented, positive-sense single-stranded RNA viruses with large (~30 kilobases) genomes that exhibit highly conserved genomic organization (Perlman and Netland, 2009; Wang, Grunewald and Perlman, 2020). While all CoVs encode 16 nonstructural proteins that function primarily in replication and transcription as well as structural proteins, each CoV subgenera encodes a unique set of interspersed accessory proteins that are dispensable for CoV replication but serve important roles in host immune evasion (Perlman and Netland, 2009; Xiang et al., 2014). For example, within the betacoronavirus genus, MERS-CoV (a merbecovirus) encodes well-characterized immune antagonists NS4a and NS4b while SARS-CoV-2 (a sarbecovirus) lacks these accessory proteins (Yang *et al*., 2013; Comar *et al*., 2019). Additionally, the nonlethal HCoVs tend to encode fewer accessory proteins than the lethal HCoVs, which may partially explain differences in pathogenesis (Fang *et al*., 2021). Despite differences among HCoVs in disease severity, relatively few studies take a comparative approach to understand CoV replication and how these viruses may interact uniquely with the host.

All successful respiratory pathogens enter and establish a primary infection in the nasal epithelium, despite its physical barrier function with apical tight junctions and robust mucociliary clearance machinery (Hiemstra, McCray and Bals, 2015; Hariri and Cohen, 2016). The nasal epithelium also serves as an important immune sentinel site where innate immune responses such as antimicrobial peptide production, interferon (IFN) production and signaling, and cytokine/chemokine signaling to recruit immune cells is initiated. Basal expression levels of IFNs and IFN-stimulated genes (ISGs) are particularly high in the nasal epithelium, suggesting that this primary barrier site may be primed for response to invading viral pathogens (Li *et al*., 2021; Loske *et al*., 2022). These barriers and host responses in the nasal epithelium likely play important roles in limiting the spread of HCoVs and other respiratory pathogens to the lower airway, preventing more severe airway disease such as lethal pneumonia. Studies comparing bronchial and nasal epithelial cells also highlight differences in innate immune responses in the upper vs. lower airway in the context of exposure to inflammatory stimuli and microbial antigens (Comer, Elborn and Ennis, 2012; Hawley *et al*., 2015). Nasal epithelial cells tended to respond more robustly to inflammatory stimuli (cigarette smoke, bacterial lipopolysaccharides) than donor-matched bronchial cells, further suggesting that the nose may be primed to respond to invading microbes (Comer, Elborn and Ennis, 2012).

Various lines of evidence highlight the importance of the nasal epithelium in CoV pathogenesis. It has been proposed that aerosolized viral particles achieve the greatest deposition density in the nasal cavity, leading to primary infection in the nose, followed by viral spread to the lower airway via a nasal/oral-lung aspiration axis (Booth *et al*., 2005; Farzal *et al*., 2019). Aspiration from the nasal/oral-pharynx occurs even in healthy individuals during sleep, and its role in other lower airway pathologies is widely recognized (Gleeson, Eggli and Maxwell, 1997; Gaeckle *et al*., 2020). Recent work with SARS-CoV-2 also emphasizes the role of nasal epithelial immune responses as determinants of pathogenicity. For example, a single-cell RNA sequencing study comparing nasopharyngeal swabs in mild and moderate COVID-19 patients found that, despite similar viral loads, patients with mild symptoms showed strong induction of antiviral IFN response genes in the nose, whereas patients with more severe symptoms had muted antiviral responses (Ziegler *et al*., 2021). Other studies have shown that early innate immune responses in the nose have a direct impact on early viral replication levels, which have been correlated with the likelihood of transmission (Cheemarla *et al*., 2021). Thus, control of HCoV replication and elimination at the initial site of infection (the nasal epithelium) is critical for the prevention of more severe symptoms and spread to the lower airway, as well as for reducing transmission (Gómez-Carballa *et al*., 2022).

We previously utilized a primary epithelial culture system in which patient-derived nasal cells are grown at an air-liquid interface (ALI) to characterize replication and induction of innate immunity by SARS-CoV-2 (Li *et al*., 2021). These nasal ALI cultures closely recapitulate many features of the *in vivo* nasal epithelium, such as cell types present (ciliated cells, mucus-producing goblet cells, as well as basal cells that repopulate the epithelium as cells senesce) and functions (epithelial barrier integrity and mucociliary function), and thus provide an optimal system in which to study HCoV replication and HCoV-host interactions (Tamashiro *et al*., 2009; Pezzulo *et al*., 2011; Lee *et al*., 2016, 2017; Kohanski *et al*., 2018). Additionally, nasal ALI cultures can be manipulated with various cytokine or drug treatments to induce changes in the airway epithelium that mirror specific disease states. For example, treatment of these cultures with IL-13, a type 2 cytokine that is known to play a role in allergy and asthma pathogenesis in humans along with IL-4 and IL-5, induces goblet cell hyperplasia and mucus hypersecretion, replicating the tissue landscape in an asthmatic airway (Ordoñez *et al*., 2001; Rogers, 2002; Kanoh, Tanabe and Rubin, 2011; Everman, Rios and Seibold, 2019). The impact of IL-13 treatment on HCoV replication is of particular interest, as most clinical association studies have shown that individuals with allergic asthma (mediated by type 2 cytokines such as IL-13) are either less prone to developing severe COVID-19 or are at no increased risk than the general population despite airway remodeling induced by asthma (Chhiba *et al*., 2020; Green *et al*., 2021; Dolby *et al*., 2022). Two recent studies treated primary bronchial epithelial cells with IL-13 and showed that this treatment resulted in significant decreases in SARS-CoV-2 replication (Bonser *et al*., 2022; Morrison *et al*., 2022). No studies have investigated the impact of type 2 immunity on MERS-CoV or HCoV-NL63 infection.

To further understand HCoV infection in the nasal epithelium, we infected donor-matched nasal ALI cultures with three HCoVs: SARS-CoV-2, MERS-CoV, and HCoV-NL63. These viruses represent both lethal and seasonal/common HCoVs, as well as alpha- and beta-coronaviruses. Notably, SARS-CoV-2 and HCoV-NL63 utilize the same cellular receptor for entry (ACE2), whereas MERS-CoV uses a different receptor (dipeptidyl peptidase 4, DPP4) (Raj *et al*., 2013; Cuervo and Grandvaux, 2020; Hoffmann *et al*., 2020; Castillo *et al*., 2022). We first characterized viral replication, the impact of temperature on replication, and host cell tropism for each virus, and then evaluated the degree to which each HCoV induced cytotoxicity in the nasal epithelium. We then extended our studies of HCoV replication and induced cytotoxicity in the nose by infecting nasal ALI cultures with HCoV-229E, (another seasonal HCoV that utilizes a different receptor, aminopeptidase N (APN) (Yeager *et al*., 1989). Finally, we treated nasal cultures with IL-13 to recapitulate an asthmatic airway and determined the impact of IL-13 on HCoV receptor abundance and replication. This comparative study seeks to further understand differences among HCoVs and how they interact with the host at the primary barrier site to infection, the nasal epithelium.

## RESULTS

### SARS-CoV-2, MERS-CoV, and HCoV-NL63 replicate productively in nasal epithelial cultures

In order to compare the replication kinetics of SARS-CoV-2, MERS-CoV, and HCoV-NL63 in the nasal epithelium, nasal air-liquid interface (ALI) cultures derived from six independent donors were infected apically at a multiplicity of infection (MOI) of 5 plaque-forming units per cell (PFU/cell) with each virus, and cultures were incubated at 33ºC to replicate the temperature of the nose *in vivo* (Keck *et al*., 2000). At 48-hour intervals following viral inoculation, apical surface liquid (ASL) was collected, and infectious virus was quantified by plaque assay on VeroE6 (for SARS-CoV-2), VeroCCL81 (for MERS-CoV), or LLCMK2 (for HCoV-NL63) cells as previously described (Schildgen *et al*., 2006; Li *et al*., 2021). No infectious virus was detected in basal media from any of the infected cultures at any time point following infection by SARS-CoV-2, MERS-CoV, or HCoV-NL63. **Figure 1A** depicts viral titer for each virus averaged from the six cultures at each time point, highlighting differences in replication kinetics among the viruses. All three CoVs replicate productively in primary nasal cells, reaching peak viral titer at 96 hours post infection (hpi). SARS-CoV-2 replicates most robustly in these cultures, reaching peak viral titers on average ten-fold higher than either MERS-CoV or HCoV-NL63. Additionally, while both SARS-CoV-2 and MERS-CoV titers plateau and remain at peak levels at the 144hpi time point, HCoV-NL63 titers decrease after 96hpi.

**Figure 1.**
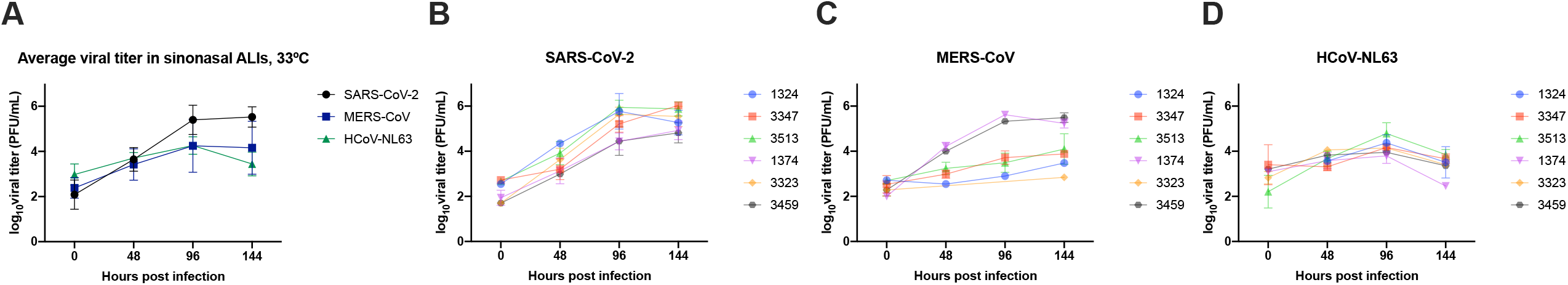
SARS-CoV-2, MERS-CoV, and HCoV-NL63 replicate productively in nasal epithelial cultures. Nasal ALI cultures derived from six donors were infected in triplicate with SARS-CoV-2, MERS-CoV, or HCoV-NL63 at MOI = 5 PFU/cell. Apical surface liquid (ASL) was collected at 48, 96, and 144 hours post infection (hpi), and infectious virus was quantified by plaque assay. (A) Titers from each of the six donors were averaged for each time point and depicted as mean ± standard deviation (SD) for each virus. (B-D) Average titers for SARS-CoV-2 (B), MERS-CoV (C), and HCoV-NL63 (D) infected cultures derived from individual donors. Donor numbers are shown in the key to the right. Each time point represents averaged titer from 3 transwells derived from that donor, displayed as mean ± SD.

In addition to these differences in average replication kinetics, within- and between-donor comparisons reveal further differences between SARS-CoV-2, MERS-CoV, and HCoV-NL63. **Figures 1B-D** depict growth curves for each virus in nasal cultures derived from each donor. Donor-to-donor variability is most evident during MERS-CoV infection (**Figure 1C)**, as it results in a bimodal replication phenotype. While nasal cultures derived from certain donors are highly susceptible to MERS-CoV infection (donors 1374, 3459), those from other donors show minimal replication of MERS-CoV. Interestingly, those donors in which MERS-CoV replicated most efficiently (donors 1374, 3459) showed the least virus production for SARS-CoV-2 (**Figure 1B**) and HCoV-NL63 (**Figure 1D**).

### Replication of SARS-CoV-2 and HCoV-NL63 but not MERS-CoV is modulated by temperature in nasal epithelial cultures

The physiologic temperature in the nasal cavity ranges from 25ºC (at the nares) to 33ºC (in the nasopharynx), which contrasts with that in the lung, which matches ambient body temperature, (37ºC) (Keck *et al*., 2000; Lindemann *et al*., 2002). Given these temperature differences in the nose, we infected nasal ALI cultures at 33ºC or 37ºC with SARS-CoV-2, MERS-CoV, and HCoV-NL63 and collected ASL as above to determine the impact of temperature on the replication of each virus (**Figure 2)**. Replication of HCoV-NL63 (at all time points) and SARS-CoV-2 (at the late time point, 144 hpi) was more efficient at 33ºC (the temperature closer to that of the *in vivo* nasal epithelium) than at 37ºC. This difference in replication was more robust for HCoV-NL63 (**Figure 2C**), as its replication was almost completely ablated when infections were conducted at 37ºC. For SARS-CoV-2 (**Figure 2A**), replication was significantly higher at 33ºC vs 37ºC only at the late time point (144hpi), consistent with another report comparing SARS-CoV-2 replication in ALI cultures derived from the lower airway (V’kovski *et al*., 2021). In contrast to SARS-CoV-2 and HCoV-NL63, MERS-CoV replication was not dependent on temperatures, as titers were not significantly different at 33ºC vs. 37ºC

**Figure 2.**
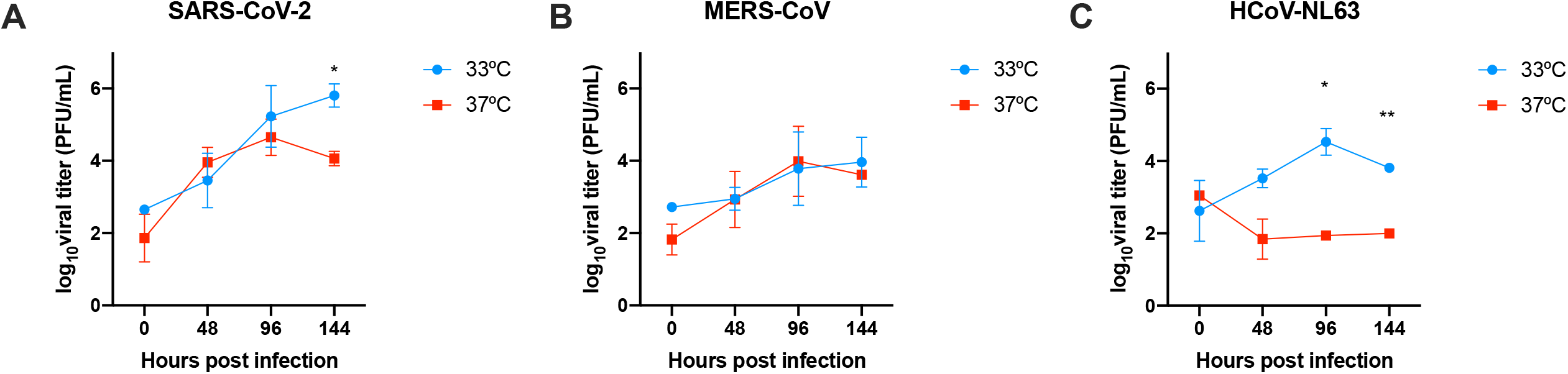
Replication of HCoV-NL63 and SARS-CoV-2, but not MERS-CoV, is modulated by temperature. Nasal ALI cultures were equilibrated at 33ºC or 37ºC for 1 day prior to infection, then were infected at MOI = 5 PFU/cell in triplicate, and incubated at 33ºC or 37ºC. ASL was collected at 48, 96, and 144 hpi and viral titers quantified via plaque assay. (A-B) Average titers from triplicate cultures derived from 7 donors infected with SARS-CoV-2 and MERS-CoV are shown as mean ± SD. (C) Average titers from triplicate cultures derived from 4 donors infected with HCoV-NL63. Statistical significance of changes in viral titer in cultures incubated at 33ºC vs. 37ºC for each virus was calculated by repeated measures two-way ANOVA: *, *P* ≤ 0.05; **, *P* ≤ 0.01. Comparisons without astericks are nonsignificant.

### SARS-CoV-2 and HCoV-NL63 infect primarily ciliated cells, while MERS-CoV infects goblet cells

After differentiation, the cell types present in the *in vivo* nasal epithelium, including ciliated cells, mucus-producing goblet cells, and basal cells that continually grow and differentiate to replace dying cells, are represented in nasal ALI cultures. We determined the cellular tropism for each CoV in the nasal epithelium using an immunofluorescence (IF) assay. Antibodies against cilia marker type IV β-tubulin and mucin MUC5AC were used to identify ciliated cells and goblet cells, respectively. Infected regions within each ALI culture were identified using antibodies against the nucleocapsid (N) protein of each CoV. Among images obtained from nasal cultures derived from 12 independent donors, both SARS-CoV-2 and HCoV-NL63 primarily infect ciliated cells, while MERS-CoV predominantly infects non-ciliated goblet cells. Representative images for each virus are depicted in **Figure 3**. This pattern is consistent with the cellular receptors used by these viruses: SARS-CoV-2 and HCoV-NL63 use the same receptor (ACE2) and thus infect the same cell type, while MERS-CoV uses DPP4.

**Figure 3.**
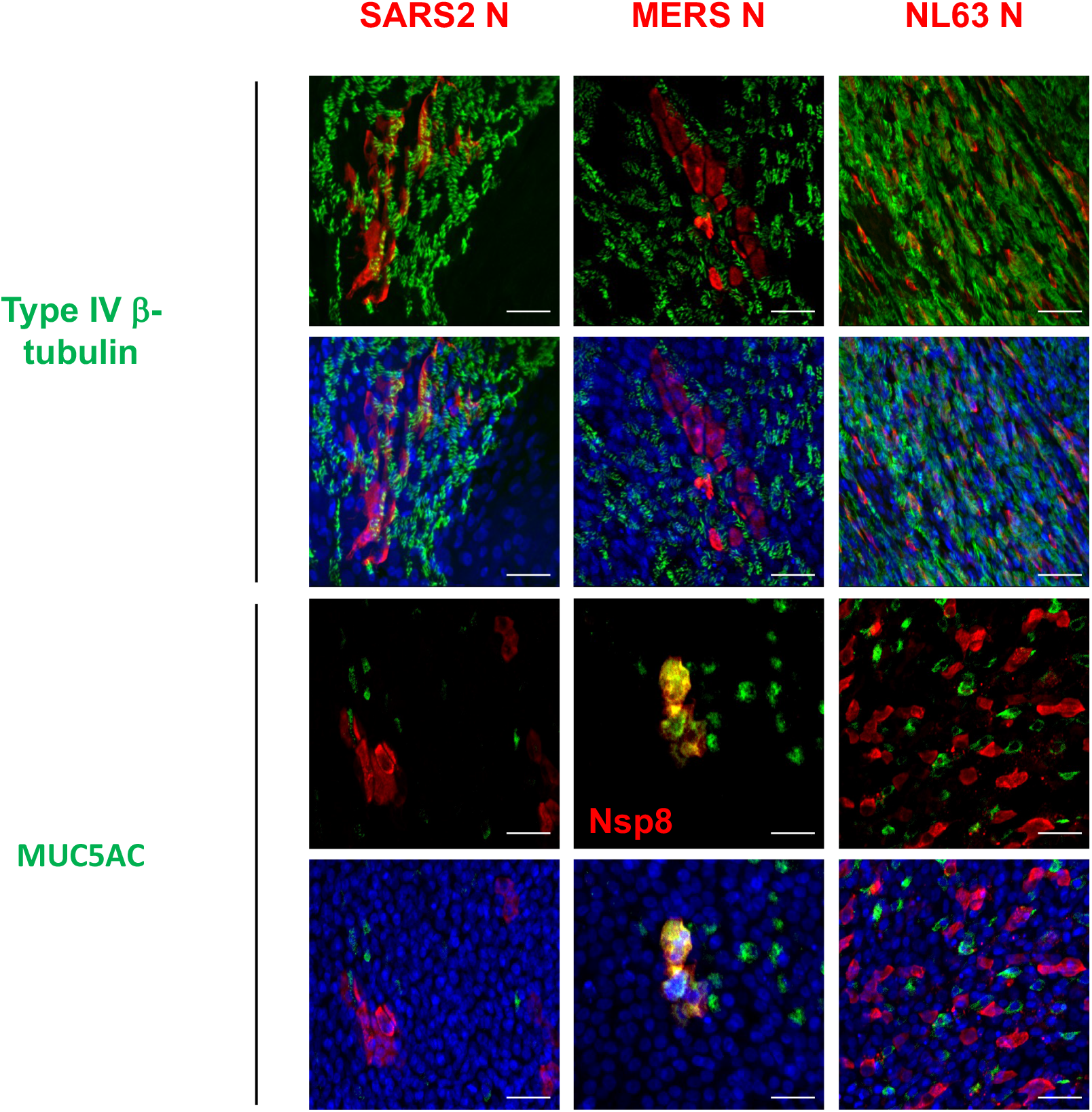
SARS-CoV-2 and HCoV-NL63 infect ciliated nasal epithelial cells, while MERS-CoV infects goblet cells. Representative images of infected nasal ALI cultures to identify cellular tropism. Infected cells were identified using primary antibodies against viral nucleocapsid for each HCoV. Primary antibodies against ciliated cell marker Type IV-tubulin and mucin MUC5AC were used to identify ciliated epithelial cells and goblet cells, respectively. Nuclei were stained with Hoescht. Note for MERS-CoV, an antibody specific to MERS-CoV nonstructural protein 8 (nsp8) was used in place of MERS-CoV nucleocapsid due to species incompatibility with the MUC5AC antibody. Images shown are representative of images acquired from nasal ALI cultures derived from 16 donors; three high-power (40X magnification) fields were analyzed in a transwell derived from each donor infected with each HCoV. Scale bars in each image are 50 μm. Representative images from mock-infected cultures stained with Type IV-tubulin, MUC5AC, and each HCoV nucleocapsid antibody confirming the absence of nonspecific signal can be found in **Supplement S1**.

### HCoVs differentially impact epithelial barrier integrity during infection of nasal epithelial cultures

To evaluate the impact of SARS-CoV-2, MERS-CoV, and HCoV-NL63 infection on epithelial barrier integrity in the nasal epithelium, nasal ALI cultures derived from 11 donors were infected with each virus, and an epithelial volt/ohm-meter (EVOM) instrument was used to measure trans-epithelial electrical resistance (TEER) in each culture prior to infection (0hpi) as well as at 96 and 192hpi (Srinivasan *et al*., 2015). Loss of epithelial integrity, tight junction dissolution, and other forms of damage to the epithelium result in decreases in TEER. To quantify global changes in epithelial barrier integrity throughout the course of infection with each virus, TEER values before infection were subtracted from TEER values at 192hpi and these values are plotted as ∆TEER in **Figure 4A**. Each point in the scatterplot represents a single transwell culture, and the superimposed bars represent the average ∆TEER for each virus among triplicate cultures derived from all 11 donors. On average, both SARS-CoV-2 and HCoV-NL63 result in negative values for ∆TEER (decreases in epithelial barrier integrity with infection), while MERS-CoV infection results in positive values for ∆TEER, similar to the increase in TEER seen in mock-infected cultures following the differentiation period. This decrement in TEER is larger in magnitude for HCoV-NL63 than SARS-CoV-2.

**Figure 4.**
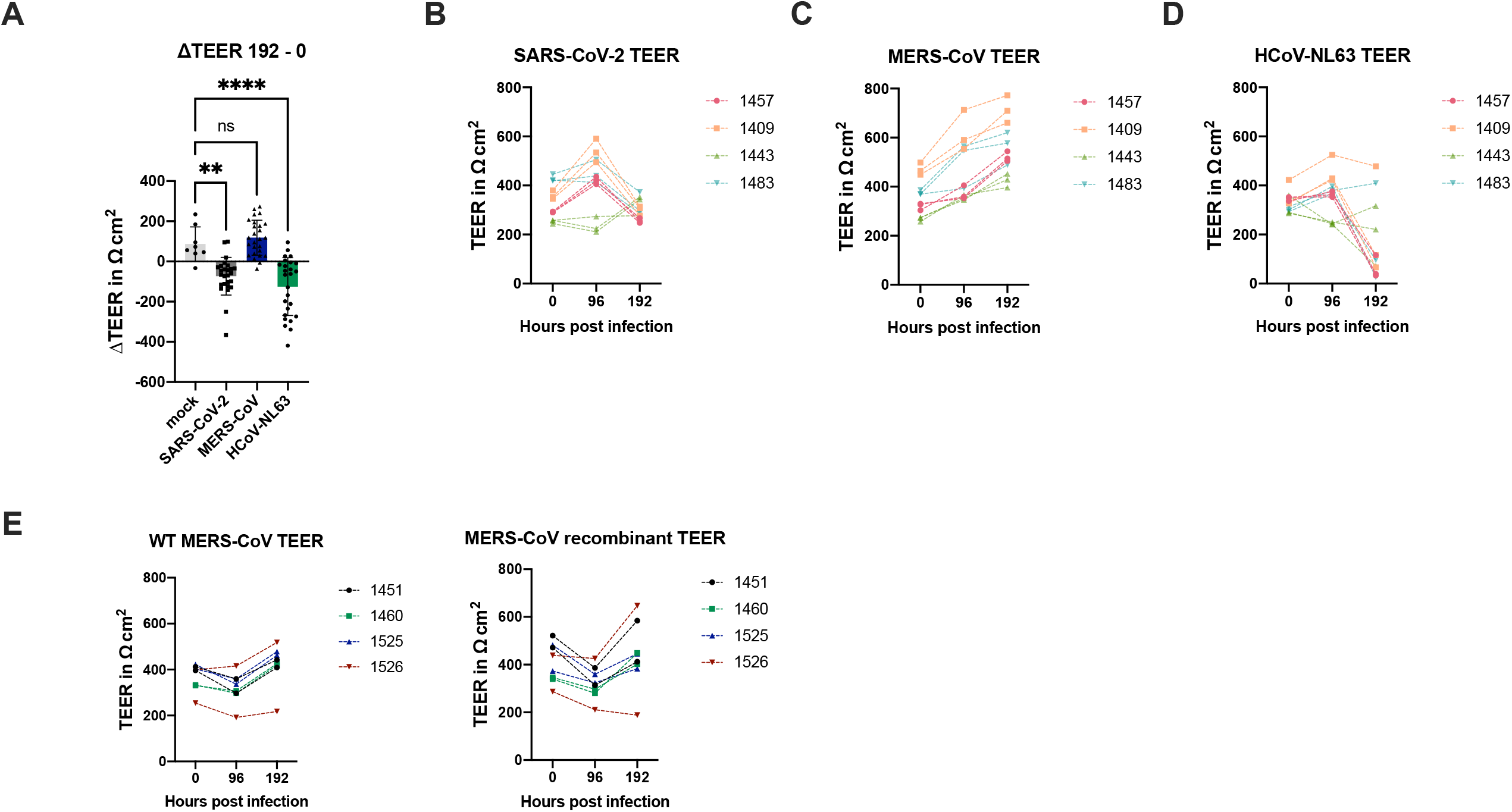
SARS-CoV-2 and HCoV-NL63 disrupt nasal epithelial barrier integrity. Nasal ALI cultures derived from 11 donors were infected with SARS-CoV-2, MERS-CoV, or HCoV-NL63 at MOI = 5 PFU/cell. Trans-epithelial electrical resistance (TEER) was measured prior to infection (0 hpi) and at 96 and 192 hpi. (A) ∆TEER values were calculated by subtracting baseline TEER (0 hpi) from TEER at 192 hpi. Each point on the scatterplot denotes the ∆TEER value for an individual transwell; data from duplicate/triplicate cultures from 11 donors is shown. The average ∆TEER is shown as a bar graph for each virus, depicted as mean ± SD. (B-D) TEER values for individual transwells derived from 4 donors infected with each HCoV are shown, illustrating TEER changes within-transwell over time. This data is representative of TEER traces acquired from cultures derived from 11 donors. (E) TEER values for individual transwells are shown for duplicate cultures derived from 4 donors and infected with WT MERS-CoV or MERS-CoV recombinant (MERS-CoV-nsp15^H231A^/∆NS4A). For TEER traces (B-E), transwells derived from the same donor are color-coded, with donor numbers shown in the key to the right. Statistical significance of ∆TEER values for each virus compared to mock-infected cultures was calculated by one-way ANOVA: **, *P* ≤ 0.01; ****, P ≤ 0.0001. Data that did not reach significance are labeled ns.

**Figures 4B-4D** depict TEER values at 0, 96, and 192hpi for nasal ALI cultures derived from 4 donors in order to monitor TEER changes in individual cultures over time. For infected cultures, TEER tends to remain stable or increase from 0hpi to 96hpi. Decreases in TEER following SARS-CoV-2 (**Figure 4B**) and HCoV-NL63 (**Figure 4D**) infection occur between 96hpi and 192hpi, while TEER continues to increase for MERS-CoV-infected cultures (**Figure 4C**). Similar TEER traces for mock-infected cultures are shown in **Figure S2**. These TEER plots highlight the donor-to-donor variability observed in nasal ALI cultures. Baseline TEER values (0hpi) and TEER values following infection tend to cluster by donor. Additionally, while SARS-CoV-2 and HCoV-NL63-infected cultures on average show a decrease in TEER over the course of infection, represented by negative ∆TEER values (**Figure 4A**), this phenotype is donor-dependent, as cultures derived from some donors that are infected with either virus have relatively minimal changes in TEER throughout infection.

To further evaluate the impact of MERS-CoV infection on epithelial barrier integrity, we evaluated TEER trends following infection with a mutant MERS-CoV recombinant virus (MERS-nsp15^H231A^/∆NS4a) that we have reported previously induces significantly stronger innate immune responses, including IFN production and signaling, as well as activation of antiviral protein kinase R and ribonuclease L, than WT MERS-CoV (**Figure 4E)** (Comar *et al*., 2022). Infection with this immune-stimulatory MERS-CoV mutant resulted in similar TEER trends as observed during WT MERS-CoV infection (increases in TEER over time). This suggests that a lack of innate immune response during MERS-CoV infection does not explain the absence of epithelial barrier destruction that is seen during SARS-CoV-2 and HCoV-NL63 infection in nasal ALI cultures.

### Infection of the nasal epithelium with SARS-CoV-2 and HCoV-NL63, but not MERS-CoV, results in significant cytotoxicity

Given that HCoV infection of nasal epithelial cultures resulted in significant changes in epithelial barrier integrity, we sought to determine whether these HCoVs resulted in detectable cytotoxicity. Nasal ALI cultures were infected with each virus, and ASL was collected at 96hpi and 192hpi for quantification of cytotoxicity via lactate dehydrogenase (LDH) release assay. LDH is an intracellular enzyme that is released upon damage to cellular membranes. To quantify cytotoxicity: LDH release from uninfected cultures (background LDH release, <2% cytotoxicity for all donors tested) was subtracted from LDH released apically from infected cultures, and this value is normalized to the LDH released from cultures treated with Triton-X 100 (maximal LDH release, 100%). These calculations reveal a phenotype that mirrors our findings for TEER (**Figure 4**). Infection with MERS-CoV results in relatively little cytotoxicity at 96hpi or 192hpi (**Figure 5A** shows individual cytotoxicity values from 4 donors). This contrasts with HCoV-NL63, which causes mild cytotoxicity at 96hpi (~20%) and significant cytotoxicity at 192hpi (~40%), as well as SARS-CoV-2, which causes minimal cytotoxicity at 96hpi but significant cytotoxicity at 192hpi (~40%) (**Figure 5A**). ASL at each time point evaluated for cytotoxicity was titered via plaque assay to confirm productive replication by all three viruses (**Figure 5B-C**). Averaged cytotoxicity values at each time point among infected cultures derived from 11 donors are plotted in **Figure 5D**. Despite productive replication by all three CoVs, MERS-CoV does not induce any significant cytotoxicity, while SARS-CoV-2 and HCoV-NL63 both induce significant cytotoxicity in the nasal epithelium. This is consistent with TEER findings, as decreases in TEER during infection were only observed following SARS-CoV-2 and HCoV-NL63 infection (concurrently with cytotoxicity). We further compared LDH release in nasal cultures infected with WT MERS-CoV and the immune-stimulatory MERS-CoV mutant to determine if the lack of cytotoxicity seen during MERS-CoV infection was related to a lack of immune response (**Figure 5E**). The MERS-CoV mutant induced a slight but nonsignificant increase in cytotoxicity than WT MERS-CoV, corroborating our TEER findings.

**Figure 5.**
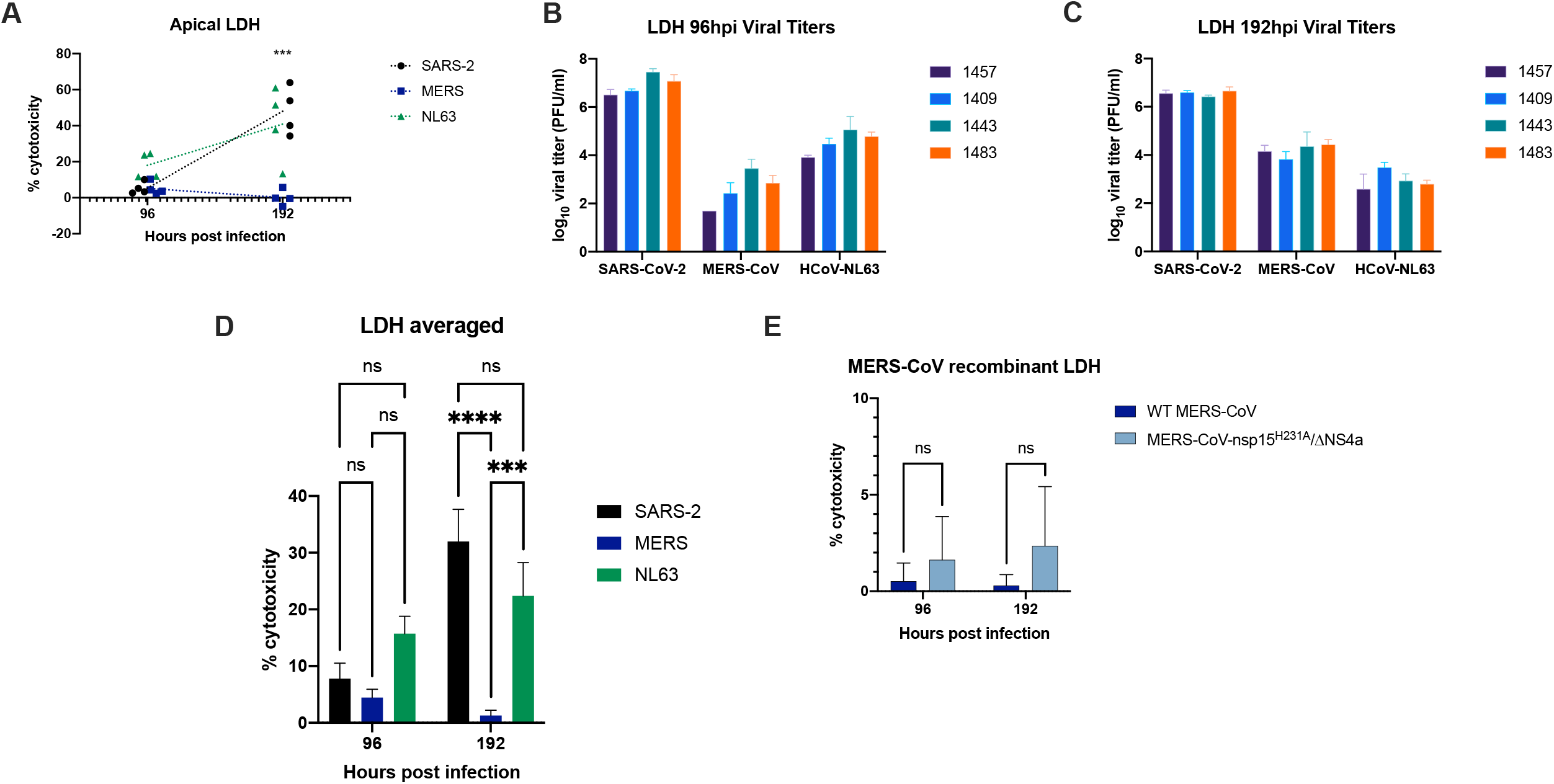
SARS-CoV-2 and HCoV-NL63, but not MERS-CoV, induce cytotoxicity in infected nasal epithelial cultures. Nasal ALI cultures derived from 11 donors were infected with SARS-CoV-2, MERS-CoV, or HCoV-NL63 and ASL was collected at 0, 96, 192 hpi. Lactate dehydrogenase (LDH) in ASL was quantified, and % cytotoxicity was calculated relative to cultures treated with 2% Triton X-100. (A) Apical LDH values from triplicate cultures derived from 4 donors infected with each HCoV were averaged and reported as mean ± SD; each point represents the average % cytotoxicity among triplicate cultures from 1 donor. Dotted lines connect the average cytotoxicity among all 4 donors for each HCoV. *** indicates average cytotoxicity in SARS-CoV-2 and HCoV-NL63 infected cultures is significantly higher than that of MERS-CoV infected cultures at 192 hpi (*P* ≤ 0.001 by two-way ANOVA). No comparisons reached significance at 96 hpi. Data from 4 donors shown here is representative of LDH data from infected cultures derived from 11 total donors. (B-C) ASL from the 4 donors in (A) was titered via viral plaque assay to confirm productive replication of each virus in these cultures. (D) % cytotoxicity values from infected cultures derived from all 11 donors were averaged and reported as mean ± SD for each virus. (E) % cytotoxicity values from duplicate cultures derived from 4 donors and infected with WT MERS-CoV or MERS-CoV recombinant (MERS-CoV-nsp15^H231A^/∆NS4A), reported as mean ± SD. Statistical significance of average % cytotoxicity values for each virus was compared at each time point via two-way ANOVA: ***, P ≤ 0.001; ****, P ≤ 0.0001. Comparisons that did not reach significance are denoted ns.

### HCoV-229E replicates productively and causes significant cytotoxicity during infection of nasal ALI cultures

To further investigate differences among lethal and seasonal HCoVs in the nasal epithelium, we infected nasal ALI cultures derived from four donors with a second alpha genus HCoV associated with the common cold, HCoV-229E. Infections were conducted at an MOI of 5 PFU/cell, and apically-shed virus was titered at 96 and 192 hpi. HCoV-229E replicates robustly in nasal ALI cultures, reaching peak titers similar to those seen for SARS-CoV-2 (**Figure 6A)**. Interestingly, HCoV-229E titers peak at 96hpi and decrease significantly at 192 hpi (~100-fold) in cultures derived from all four donors. This decline in viral titers at late time points is also observed during HCoV-NL63 infection, but not for SARS-CoV-2 or MERS-CoV (**Figure 5B-C**). We measured TEER in these cultures at 96 and 192hpi. TEER values decreased, indicating damage to epithelial barrier integrity, in three of the four donors tested between 0 and 96hpi (**Figure 6B**). Interestingly, between 96 and 192hpi, TEER values increased in all four donors. This pattern is unique among the HCoVs evaluated in this study, as the defect in TEER in the majority of donors occurred earlier during infection (between 0 and 96hpi). TEER values for SARS-CoV-2 and HCoV-NL63 decreased most significantly between 96 and 192hpi (**Figure 4B, 4D**). ∆TEER values for HCoV-229E-infected cultures as well as mock-infected cultures are plotted in **Figure 6C**, highlighting that changes in TEER during HCoV-229E infection occurred primarily between 0 and 96hpi. Finally, we measured cytotoxicity during infection with HCoV-229E via LDH quantification in ASL. Similar to the pattern seen for TEER, quantification, HCoV-229E induced a significant cytotoxicity signature (% cytotoxicity ranging from 10-40%, depending on the donor) at 96hpi (**Figure 6D**). Later in infection at 192hpi, no detectable cytotoxicity was found following HCoV-229E infection. This cytotoxicity signature is consistent with the titers seen during HCoV-229E infection, as high viral titers at 96 hpi result in negative ∆TEER values and significant LDH release into ASL, whereas lower viral titers at 192 hpi are not associated with any defect in TEER or significant LDH signature. HCoV-229E infection of nasal ALI cultures results in productive replication and evidence of cytotoxicity (decrease in TEER and LDH detection in ASL) during infection.

**Figure 6.**
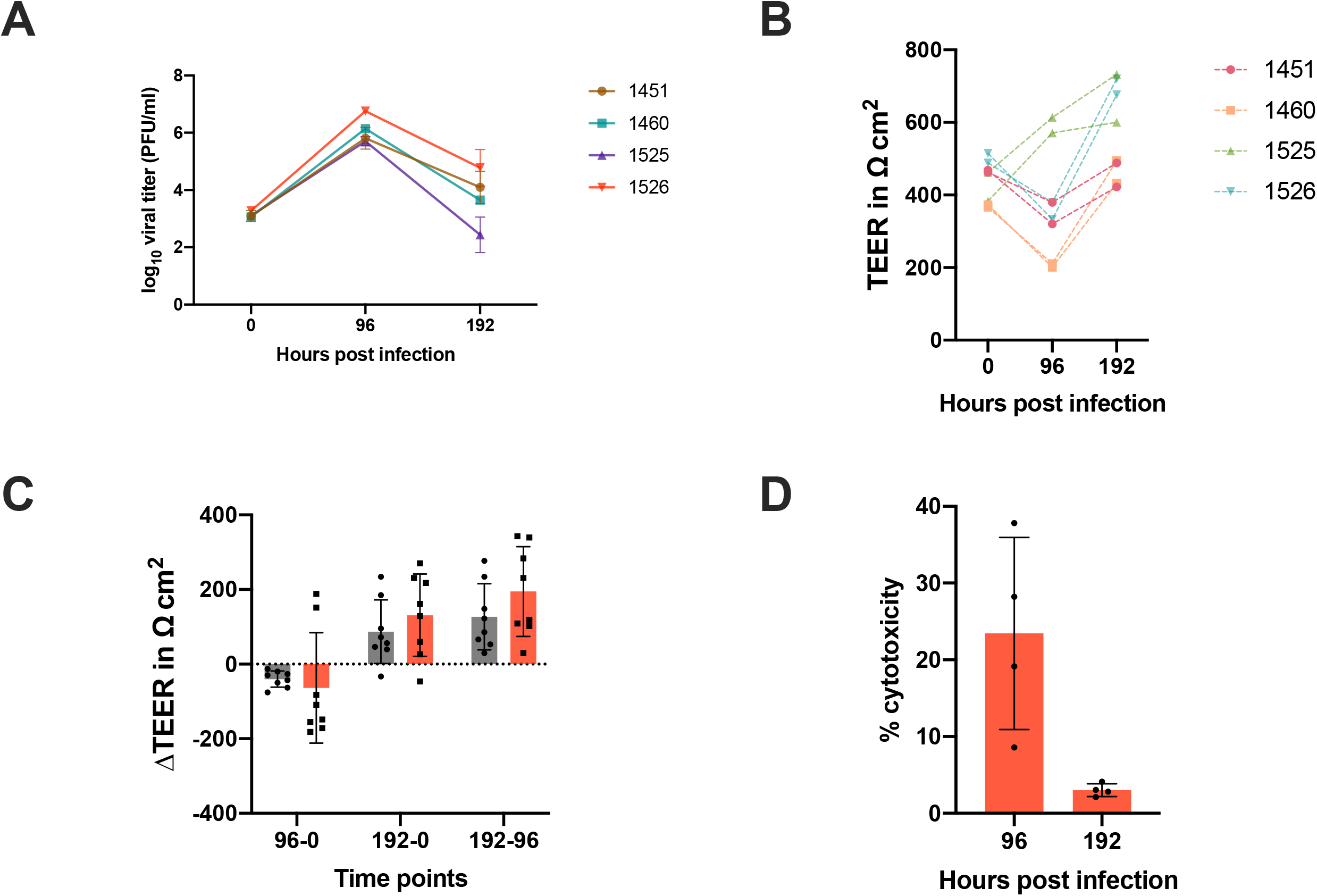
HCoV-229E replicates in nasal cultures and induces an early cytotoxicity signature. Nasal ALI cultures derived from 4 donors were infected in duplicate with HCoV-229E at MOI = 5 PFU/cell and ASL collected at 0, 96, and 192 hpi. (A) HCoV-229E viral titers determined via plaque assay, reported as mean ± SD for each of the 4 donors. (B) TEER measured prior to infection (0 hpi), as well as at 96 and 192 hpi, is plotted for each transwell infected with HCoV-229E to track TEER changes over time. Transwells derived from the same donor are color-coded according to the donor number key to the right. (C) ∆TEER values for each HCoV-229E infected culture were calculated by subtracting TEER at baseline from 96hpi TEER values (96-0), subtracting TEER at baseline from 192hpi TEER values (192-0), or by subtracting 96 hpi TEER values from 192 hpi TEER values (192-96). Each point represents ∆TEER calculated from a single transwell, with bars indicating average ∆TEER for that time point comparison. (D) ASL was used for LDH quantification to determine % cytotoxicity relative to nasal cultures treated with Triton X-100 (ceiling). Each point represents the average % cytotoxicity from duplicate transwells from each donor, and the overall average % cytotoxicity is shown with bars.

### IL-13 robustly influences cellular distribution and HCoV receptor abundance in the nasal epithelium

IL-13 is a type 2 cytokine implicated in allergy and asthma pathogenesis which causes marked goblet cell hyperplasia *in vivo* and in various ALI culture systems (Ordoñez *et al*., 2001; Atherton *et al*., 2003; Kanoh, Tanabe and Rubin, 2011). We sought to determine how IL-13 impacts cellular makeup and HCoV receptor distribution in nasal ALI cultures. To do this, we compared cultures for which differentiation media had been supplemented with IL-13 every 48 hours during the final two weeks of differentiation (IL-13 condition) vs sham-treated cultures. Using RT-qPCR with primers specific for these viruses’ cellular receptors (DPP4 and ACE2), we found that *DPP4* mRNA expression levels increased between 100- and 1000-fold relative to sham-treated cultures (**Figure 7A**), while expression of *ACE2* mRNA did not change significantly with IL-13 treatment (**Figure 7B**). We next confirmed that IL-13 treatment of nasal ALI cultures resulted in goblet cell hyperplasia and that observed changes in RNA expression of *DPP4* were associated with increased DPP4 receptor abundance using an IF assay with antibodies against MUC5AC (goblet cell marker) and DPP4. Imaging of nasal ALI cultures derived from 10 total donors treated with IL-13 revealed marked increases in the numbers of cells positive for both MUC5AC and DPP4 via imaging (**Figure 7C**). DPP4 signal by immunofluorescence is very low at baseline (sham treatment). To confirm these findings, we collected protein lysates from mock-infected sham- and IL-13-treated cultures derived from 4 pooled donors, as well as from SARS-CoV-2, MERS-CoV, and HCoV-NL63-infected cultures at 96hpi. Consistent with *DPP4* mRNA expression, DPP4 protein levels were undetectable by western blot in sham-treated cultures but present in all IL-13-treated cultures (**Figure 7D**). Though *ACE2* mRNA expression was not impacted by IL-13 treatment (**Figure 7B**), ACE2 protein levels decreased with IL-13 treatment (**Figure 7D**), consistent with a previous study (Kimura *et al*., 2020). We also evaluated total type IV β-tubulin (ciliated cell marker) and MUC5AC levels (goblet cell marker), which showed reciprocal patterns with IL-13 treatment. MUC5AC protein levels increased with IL-13 treatment (further confirming goblet cell hyperplasia), while type IV β-tubulin protein levels decreased (suggesting that ciliated cells may diminish in number secondary to IL-13 treatment). These data demonstrate that IL-13 treatment has significant effects on cellular distribution (goblet vs ciliated cells) and receptor expression level (DPP4 for MERS-CoV, ACE2 for SARS-CoV-2 and HCoV-NL63) in nasal ALI cultures.

**Figure 7.**
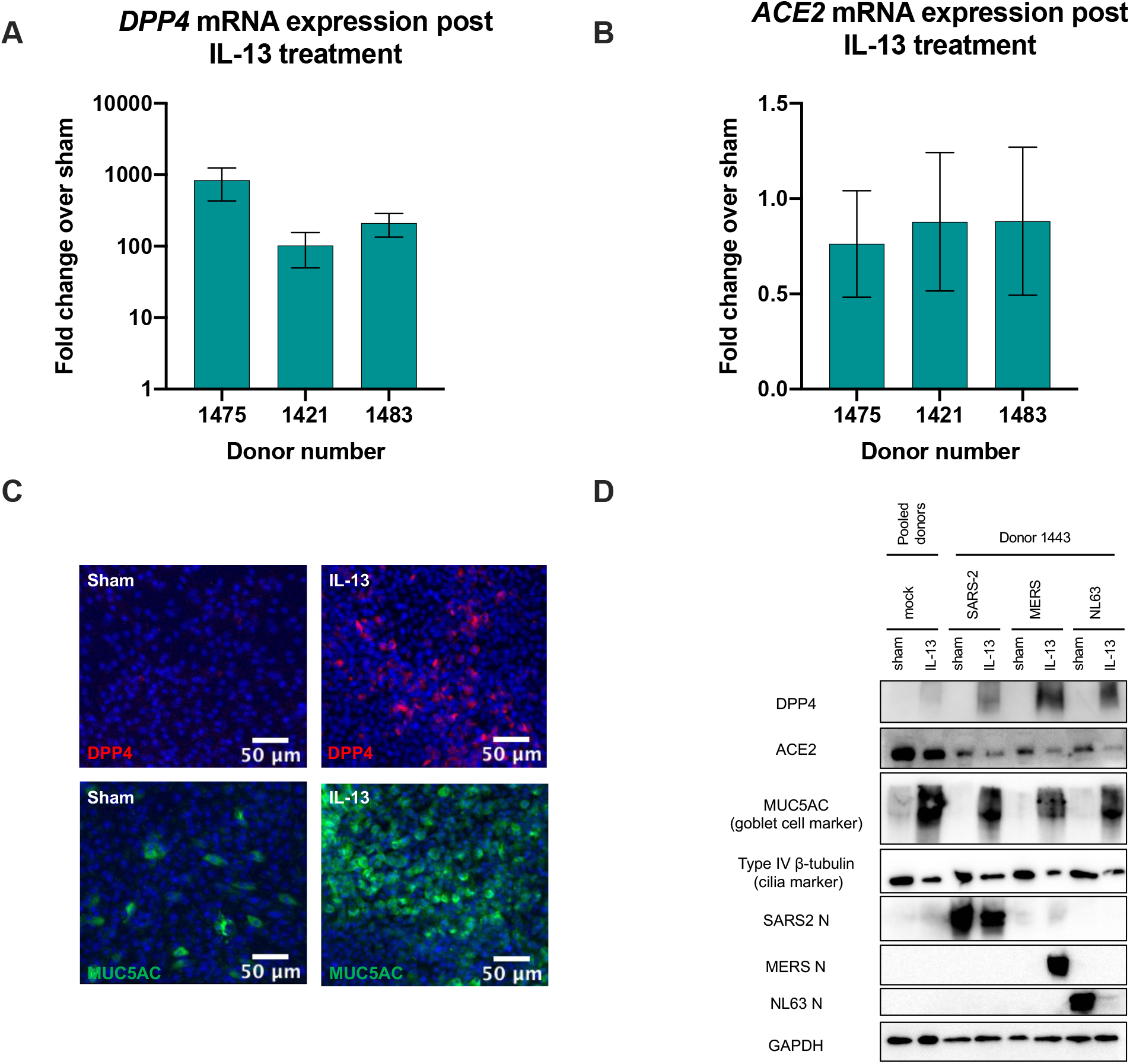
IL-13 treatment of nasal epithelial cultures impacts HCoV receptor abundance and cellular distribution. (A-B) Uninfected nasal ALI cultures derived from 3 donors were sham-treated or treated with IL-13 in triplicate for the final 2 weeks of differentiation, and total RNA was harvested and expression of *DPP4* (A) and *ACE2* (B) mRNA was quantified by RT-qPCR and expressed as fold change over sham-treated cultures using the 2^−Δ(Δ*CT*)^ formula. Data are displayed as means ± SD. (C) Uninfected sham- and IL-13-treated ALI cultures were fixed and stained with primary antibodies against DPP4 and MUC5AC. Representative images from 1 of 4 donors analyzed in this way are shown. (D) Uninfected ALI cultures derived from pooled donors (cells from 4 donors pooled prior to plating on transwells) were sham- or IL-13-treated and total protein was harvested for analysis via western blot. ALI cultures derived from 10 donors (for SARS-CoV-2 and MERS-CoV) or 7 donors (for HCoV-NL63) were sham- or IL-13-treated prior to infection with each HCoV and total protein harvested for similar western blot analysis. Proteins were separated via SDS-PAGE and immunoblotted with antibodies against DPP4, ACE2, MUC5AC, Type IV-tubulin, Nucleocapsid (N) for each of SARS-CoV-2 (SARS-2), MERS-CoV (MERS), and HCoV-NL63 (NL63), and GAPDH. Data are representative of similar findings in 10 (for SARS-CoV-2 or MERS-CoV) or 7 (for HCoV-NL63) donors total.

### SARS-CoV-2 and HCoV-NL63 replication is inhibited, while MERS-CoV replication increases, following IL-13 treatment of nasal ALI cultures

We next aimed to evaluate how IL-13 treatment and the resulting changes in HCoV receptor availability and cellular distribution impacted the replication of SARS-CoV-2, MERS-CoV, and HCoV-NL63. To do this, we treated donor-matched nasal ALI cultures derived from 10 donors (for SARS-CoV-2 and MERS-CoV) or 7 donors (for HCoV-NL63) with IL-13 during the final two weeks of differentiation as above, infected at an MOI of 5 PFU/cell with each virus, and collected ASL at 48hpi and 96hpi for titration via plaque assay. We observed dramatic changes in the replication of all three HCoVs with IL-13 treatment (vs. replication in sham-treated cultures). Donor-matched before-and-after plots are shown in **Figure 8**, depicting average viral titer at 48 (**8A**) and 96 hpi (**8B)** for each donor in sham vs. IL-13-treated cultures, connected with a line. **Figure S3** depicts average viral titer among all donors in sham- vs. IL-13-treated cultures. On average, both SARS-CoV-2 and HCoV-NL63 replicate less efficiently in IL-13-treated vs. sham-treated cultures. This phenotype is consistent with the cellular tropism and receptor expression for these viruses, as IL-13 treatment resulted in decreased ACE2 expression as well as decreased IV β-tubulin levels at the protein level via western blot (**Figure 7D**). Interestingly, HCoV-NL63 replication decreased more dramatically than that of SARS-CoV-2 following IL-13 treatment, despite utilizing the same cellular receptor. MERS-CoV replication, on average, increased with IL-13 treatment, consistent with its goblet cell tropism and use of the DPP4 receptor, which both increased in abundance after IL-13 treatment (**Figure 7B-D**). These differences in viral replication following IL-13 treatment occur early in infection (48 hpi) and are sustained later in infection (96 hpi). The before-and-after plots in **Figure 8A-B** highlight the donor-to-donor variability that we observe throughout our studies. For SARS-CoV-2 and MERS-CoV, there is some donor-dependent variation in how dramatically IL-13 treatment impacts each virus’ replication.

**Figure 8.**
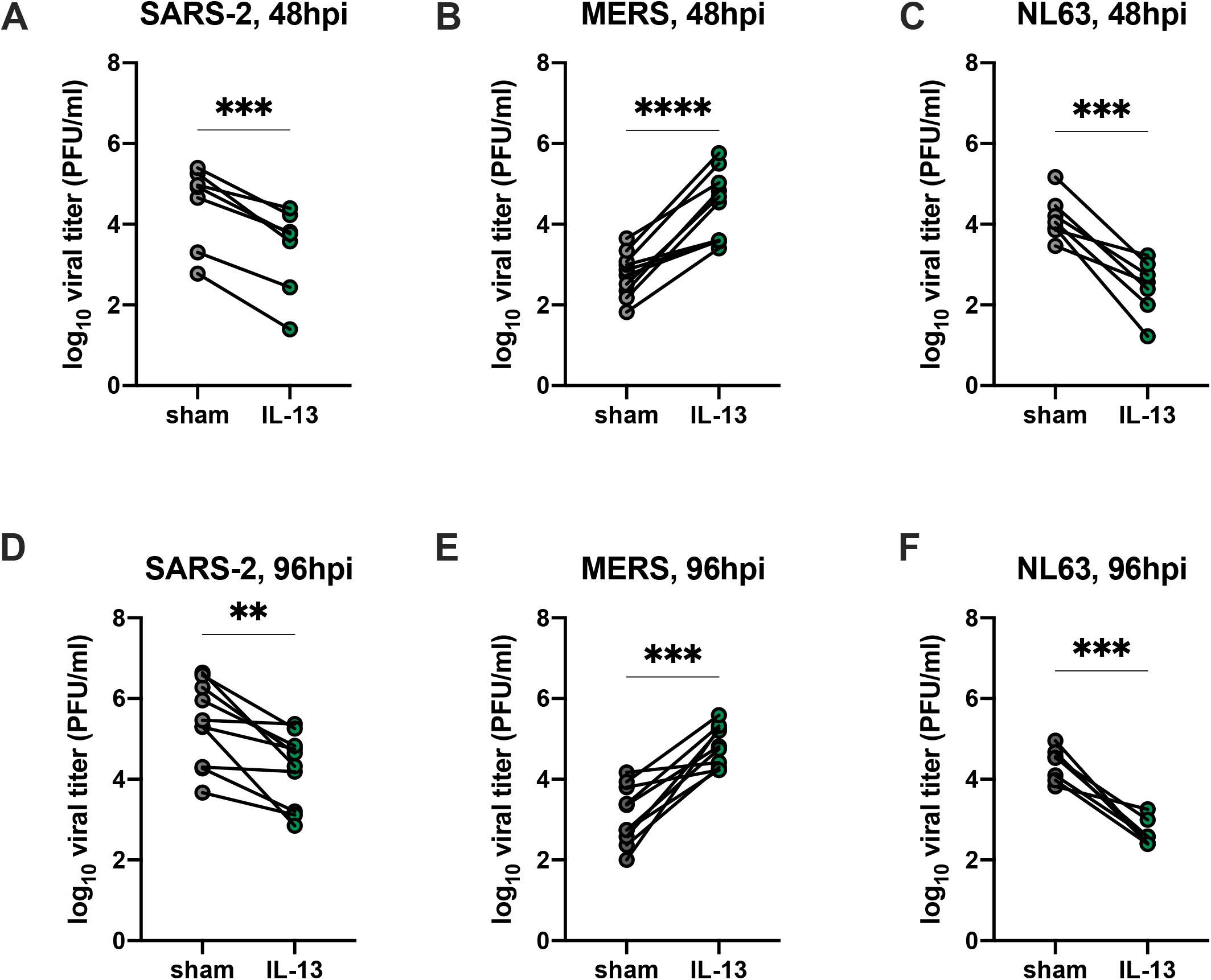
HCoV replication is significantly impacted by IL-13 treatment of nasal epithelial cultures. Nasal ALI cultures derived from 10 donors (for SARS-CoV-2 and MERS-CoV) or 7 donors (for HCoV-NL63) were sham- or IL-13-treated for the final 2 weeks of differentiation and then were infected in triplicate with each HCoV. ASL was collected at 48 and 96 hpi and infectious virus quantified via plaque assay. (A-C) Viral titer at 48 hpi for sham- and IL-13-treated cultures infected with each HCoV was averaged by donor and plotted as before-and-after plots, connecting mean titer in sham-treated cultures (gray) with mean titer in IL-13-treated cultures (green). Each set of connected lines represents titer data from 1 donor. (D-E) Similar before-and-after plots comparing 96 hpi titers in donor-matched sham- vs. IL-13 treated cultures. Statistical significance of the difference in titer in sham vs. IL-13 treated cultures was determined for each HCoV at each time point using paired t-tests: **, *P* ≤ 0.01; ***, P ≤ 0.001; ****, P ≤ 0.0001.

## DISCUSSION

We have utilized a primary culture system in which patient-derived nasal epithelial cells are grown at an air-liquid interface (ALI) to identify differences among pathogenic (SARS-CoV-2 and MERS-CoV) and seasonal (HCoV-NL63 and HCoV-229E) HCoVs. This nasal ALI system offers many advantages over the use of traditional epithelial cell lines to study HCoV replication. First, the use of differentiated nasal epithelial cells allows us to study virus-host dynamics at the primary barrier site encountered by respiratory viruses. Comparative studies of upper vs. lower respiratory tract cell lines (as well as primary culture systems) identify differences in permissiveness to viral infection as well as in host responses and transcriptional profiles, highlighting the need to study HCoVs in nasal cell models. Relatively few nasal epithelial cell lines are available, and the few that are (such as RPMI 2650, used as a model for chronic rhinosinusitis) have been shown to have limited response to inflammatory stimuli compared to primary nasal cells (Ball *et al*., 2015). Additionally, nasal ALIs express the cell types and mucociliary functions present in the nose, recreating the tissue environment and primary barriers encountered by HCoVs in the airway. Many groups have also shown that ALI systems like this one replicate the transcriptional profile of *in vivo* airway epithelia more closely than submerged culture conditions (Pezzulo *et al*., 2011).

Prior studies have demonstrated that primary nasal epithelial cells are highly susceptible to SARS-CoV-2 infection, which is at least partially explained by the relatively high expression levels of ACE2 in the upper respiratory tract (Hou *et al*., 2020; Hatton *et al*., 2021; Gamage *et al*., 2022; Tran *et al*., 2022). Notably, relatively few studies have taken a comparative approach to understand the diversity among HCoVs in their replication and concurrent host responses, and our work is the first to rigorously compare SARS-CoV-2, MERS-CoV, HCoV-NL63, and HCoV-229E in a nasal epithelial culture system. Most studies on MERS-CoV have focused on its replication in the lower respiratory tract, given its propensity to cause severe pneumonia; however, our data using nasal ALI cultures have demonstrated that MERS-CoV can productively replicate in the nasal epithelium, and, presumably, infection with MERS-CoV begins in the nose (de Wit *et al*., 2013; Li *et al*., 2021). HCoV-NL63 and HCoV-229E, as well as the other HCoVs typically associated with the common cold, have been remarkably under-studied. However, HCoV-NL63 is known to use the same cellular receptor as SARS-CoV-2 (ACE2), and infection of bronchial/tracheal epithelial cells has shown that HCoV-NL63 replication is correlated with ACE2 expression (Castillo *et al*., 2022). Since all HCoVs enter and likely establish primary infections in the nasal epithelium, we hypothesized that understanding the diversity in HCoV-host dynamics in the nose may inform differences in disease phenotype as well as transmissibility among these viruses.

The main focus of our study was to directly compare SARS-CoV-2, MERS-CoV, and HCoV-NL63. Titration of apically shed virus in donor-matched nasal epithelial cultures revealed that, while all three HCoVs replicate productively in the nasal epithelium, SARS-CoV-2 reaches the highest viral titers. This correlates with and may explain the higher transmissibility of SARS-CoV-2 compared to MERS-CoV, in which person-to-person transmission is far less common and has only been observed in settings such as hospital outbreaks and within households. As in other ALI culture systems, we observed some donor-to-donor variability in viral replication in our cultures. This was most evident during infection with MERS-CoV, which revealed a bimodal phenotype in which some donors were much more permissive to MERS-CoV replication than others. We hypothesize that this may be related to baseline variability in cell type composition and receptor expression in nasal cultures – i.e. cultures with an increased proportion of goblet cells and increased expression of MERS-CoV receptor (DPP4) are likely more permissive to infection by MERS-CoV. This donor-to-donor variability is another advantage of this primary culture system over immortalized cell lines, as HCoVs are often associated with a spectrum of clinical disease which is likely at least partially explained by heterogeneity in susceptibility and host responses.

Given the lower temperature typically found in the nasal airway (30-35ºC rather than 37ºC in the lower airway), we compared viral replication in nasal ALI cultures incubated at 33ºC and 37ºC (Keck *et al*., 2000; Lindemann *et al*., 2002). Temperature significantly impacted HCoV-NL63, and to some extent SARS-CoV-2, but did not impact MERS-CoV replication. Temperature-dependent differences in viral replication could be explained by differences in virion stability, viral replication efficiency, or host responses. It has been reported in the context of rhinovirus and various arbovirus infections that host innate immune responses are dampened at lower temperatures, suggesting that cooler temperatures may enable viral replication (Foxman *et al*., 2015; Lane *et al*., 2018). Biochemical studies on influenza virus have revealed that temperature can impact the stability of viral replication machinery (Dalton *et al*., 2006). We are currently investigating the role of each of these factors in mediating the temperature-dependent differences in viral replication observed in nasal epithelial cultures.

To evaluate the degree of cytotoxicity induced during infection of the nasal epithelium with each HCoV, we quantified TEER as a readout for epithelial barrier integrity and LDH release as a marker for cellular damage. Decreases in TEER have been documented to occur secondary to various respiratory viral infections, including influenza and respiratory syncytial virus (Smallcombe et al., 2019; Ruan et al., 2022). While both SARS-CoV-2 and HCoV-NL63 induced marked cytotoxicity in nasal ALI cultures, resulting in negative ∆TEER values (deterioration in epithelial barrier integrity) and significant LDH signal, MERS-CoV did not induce significant cytotoxicity. Interestingly, MERS-CoV infection is not associated with significant upper respiratory tract symptomology, causing primarily severe lower respiratory tract disease (lethal pneumonia in more than 35% of cases). This contrasts with HCoV-NL63, typically associated with the common cold and primarily upper respiratory symptoms, as well as SARS-CoV-2, which is known to cause a wide range of disease from asymptomatic infections to mild colds to severe pneumonia. It is plausible that HCoV-mediated cytotoxicity in the case of mild SARS-CoV-2 and HCoV-NL63 infections in the nasal epithelium may facilitate early viral clearance and limit the subsequent spread of viral infection to the lower airway (Ramasamy, 2022). Whereas limited cytotoxicity during MERS-CoV infection may allow for uninhibited spread to cause lower airway pathology.

Given the striking differences in cytotoxicity profiles observed during infection of nasal ALI cultures among these three HCoVs, we extended these studies by including another common cold-associated HCoV (HCoV-229E). HCoV-229E replicates robustly in the nasal epithelium, reaching peak titers similar to those seen for SARS-CoV-2, but its overall replication kinetics differ, as a sharp decline in apically shed virus is observed at late time points. This decline in viral titer is also observed for HCoV-NL63 at very late time points, suggesting it may be a common feature during infection of nasal epithelial cells with seasonal HCoVs. HCoV-229E induced cytotoxicity during infection of nasal ALI cultures – indicated by negative ∆TEER values and LDH detected in apical fluid – but this cytotoxicity signature was completely resolved at the very late time point (192hpi). This diverged from the pattern seen for SARS-CoV-2 and HCoV-NL63, which both induced increasing cytotoxicity as infection progressed. The cytotoxicity pattern for HCoV-229E is consistent with its replication cycle, as peak viral titers occur simultaneously with both markers of cytotoxicity, and the decline in viral titer later in infection is accompanied by resolution of cytotoxicity. We speculate that this pattern may indicate clearance or resolution of infection with HCoV-229E, allowing infected nasal ALI cultures to return to a baseline or healthy state. The overall cytotoxicity profile for HCoV-229E is also consistent with its typical clinical phenotype, as it is associated with upper respiratory tract pathology and symptoms. Thus, we expect to see markers of cytotoxicity during infection of nasal ALI cultures with HCoV-229E, as was observed during SARS-CoV-2 and HCoV-NL63 infection. However, the kinetics of cytotoxicity induction by HCoV-229E are unique, suggesting additional differences among these HCoVs in terms of host responses and resolution of infection by the host.

Cytotoxicity during HCoV infection of the nasal airway is likely induced both by cellular remodeling as a byproduct of viral replication and by host immune/stress responses secondary to viral sensing. Nasal epithelial cells express high basal levels of antiviral interferon (IFN) response genes and thus may be primed for response to invading viruses (Hatton *et al*., 2021; Li *et al*., 2021). We previously showed that SARS-CoV-2 infection induces mild IFN responses in nasal ALI cultures, whereas MERS-CoV adeptly shuts down this host innate immune pathway unless its immune antagonist Endoribonuclease U (nsp15) and subgenera-specific accessory genes are mutated (Li *et al*., 2021; Comar *et al*., 2022). The degree of innate immune induction during HCoV-NL63 or HCoV-229E infection is unknown. Thus, future work in nasal ALI cultures will investigate the role that innate immune induction by HCoVs may have on their contrasting cytotoxicity profiles. There is evidence that early immune responses can mediate control of viral infections and prevention of spread to the lower airway. The host factor IFN-lambda is thought to play a particularly important role in these early defenses, as infection of mice defective in IFN-lambda with influenza results in an inability to control infection in the upper airway and dissemination to the lower airway (Klinkhammer *et al*., 2018). The protective role of IFN-lambda has just begun to be studied during HCoV infection (Chong *et al*., 2022). A recent study investigated potential mechanisms for the relative protection against developing severe COVID-19 seen in children compared to adults and revealed that pediatric airways showed higher basal expression levels of relevant immune sensors such as MDA5, which detects CoV double-stranded RNA (Loske *et al*., 2022). Furthermore, our observations of HCoV-229E suggest that even among the seasonal HCoVs, there may be significant differences in the ability of host responses to resolve viral infections. Differences in early cytotoxicity secondary to viral replication as well as degree of immune evasion in the nasal epithelium may be key factors that determine disease severity.

Finally, we treated nasal ALI cultures with type 2 cytokine IL-13 to induce goblet cell hyperplasia and mucus hypersecretion. Type 2 cytokines are known to play important roles in asthma and allergy pathogenesis, and recent clinical association studies have shown that asthmatics may be less prone to developing severe COVID-19. Our findings highlight how host factors such as baseline tissue microenvironment (i.e. goblet cell hyperplasia) can have significant impacts on HCoV receptor availability and resulting HCoV replication after infection. IL-13 treatment significantly increased MERS-CoV replication but decreased SARS-CoV-2 and HCoV-NL63 replication. Our findings are consistent with two recent reports that investigated the impact of IL-13 on SARS-CoV-2 replication in a lower airway ALI culture system (Bonser *et al*., 2022; Morrison *et al*., 2022). Our findings are also consistent with the clinical association studies that suggest asthmatic patients are resistant to severe COVID-19, likely secondary to baseline goblet cell hyperplasia, reduction in ACE2 expression, and diminished SARS-CoV-2 replication (Green *et al*., 2021). We hypothesize that asthmatic patients would be highly susceptible to severe MERS-CoV infection, as MERS-CoV primarily infects goblet cells, and asthmatic airways typically express higher than normal levels of DPP4 (Raj *et al*., 2013; Shiobara *et al*., 2016; Zhang *et al*., 2021). The opposite is likely true for HCoV-NL63, since ACE2 levels have been shown to be a key predictor of susceptibility to HCoV-NL63 infection, so asthmatic patients are likely at reduced risk of contracting HCoV-NL63 (Castillo *et al*., 2022). This nasal ALI culture system can thus be used as a tool to mimic human disease phenotypes and determine the impact of a disease on HCoV-host dynamics. In addition to IL-13 treatment to recapitulate an asthmatic/allergic airway, ALI cultures could be grown from pediatric and adult patients to further understand differences in HCoV replication and immune responses related to age. Similarly, ALI cultures derived from patients with respiratory diseases like cystic fibrosis could be used to determine relative risk profiles for these patients for these and future emerging respiratory viruses.

Given the ongoing nature of the COVID-19 pandemic, as well as the likelihood of emergence of additional pathogenic HCoVs in the future, it is imperative to understand CoV-host interactions to inform the development of effective antivirals and vaccines. Comparative studies using primary airway culture systems are particularly important to reveal diversity among HCoVs in terms of replication and host responses which are likely key predictors in pathogenesis and transmissibility. We are currently extending our work with this nasal ALI culture system to expand our findings to other HCoVs associated with the common cold, as well as to SARS-CoV-2 variants of concern.

## MATERIALS & METHODS

### Nasal air-liquid interface (ALI) cultures

Nasal mucosal specimens were obtained via cytologic brushing of patients in the Department of Otorhinolaryngology-Head and Neck Surgery, Division of Rhinology at the University of Pennsylvania and the Philadelphia Veteran Affairs Medical Center after obtaining informed consent. Acquisition and use of nasal specimens was approved by the University of Pennnsylvania Institutional Review Board (protocol #800614) and the Philadelphia VA Institutional Review Board (protocol #00781). Patients with history of systemic disease or on immunosuppressive medications are excluded. ALI cultures were grown and differentiated on 0.4 μm pore transwell inserts as previously described (Lee *et al*., 2016, 2017; Patel *et al*., 2019). In brief: cytologic brush specimens are dissociated and fibroblast cell population removed, followed by plating onto transwell inserts. Nasal cells are allowed to grow to confluence in submerged state (~5 days), then apical growth medium is removed. Basal differentiation media is replaced bi-weekly for 3-4 weeks prior to infection. All cultures are subjected to confirmatory tests for differentiation prior to infection: epithelial morphology monitored via microscopy and ciliation confirmed. Due to supply chain issues, two kinds of ALI culture differentiation medium were used. A 1:1 mixture of Bronchial Epithelial Cell Basal Medium (Lonza) with Dulbecco’s Modified Eagle Medium (DMEM) was used for experiments in Figures 1-3. PneumaCult-ALI basal medium (Stemcell Technologies) was used for experiments in Figures 4-8.

### Virus stocks

SARS-CoV-2 (USA-WA1/2020 strain) obtained via BEI resources was propagated in Vero-E6 cells. MERS-CoV was derived from a bacterial artificial chromosome encoding the full-length MERS-CoV genome (HCoV-EMC/2012) and was propagated in Vero-CCL81 cells. HCoV-NL63 was propagated in LLCMK2 cells. Low MOI (0.01) infections were used to generate virus stocks for all three viruses. HCoV-NL63 virus stock underwent ultracentrifugation through a 20% sucrose gradient to concentrate virus stock prior to infections.

### Infections and quantification of apically shed virus

All infections were conducted at MOI = 5 PFU/cell. Viruses were diluted in serum-free Dulbecco’s modified Eagle’s medium (DMEM) to achieve a total inoculum volume of 50 μL and added apically to nasal ALI cultures for adsorption for 1 hour. After viral adsorption, cells were washed three times with phosphate-buffered saline (PBS). At indicated time points, 200 μL PBS was added to the apical surface of each infected transwell and collected for subsequent quantification of infectious virus via standard plaque assay. Different cell lines and incubation periods were used for titration of each virus: VeroE6 for 3 days at 37ºC (SARS-CoV-2), VeroCCL81 for 4 days at 37ºC (MERS-CoV), and LLCMK2 for 6 days at 33ºC (HCoV-NL63). All virus manipulations and infections were conducted in a biosafety level 3 (BSL-3) facility using appropriate and approved personal protective equipment and protocols.

### IL-13 treatment

Basal media was supplemented with 50 ng/μL IL-13 (R&D Systems, cat # 213-ILB-010) and replaced every 48 hours for the final two weeks of differentiation before infection for IL-13-treated cultures. Sham-treated cultures were treated in the same way with Hank’s Balanced Salt Solution (Gibco, cat # 14175-079).

### Immunofluorescence (IF) staining

Following infection, the cultures were washed 3 times with PBS and fixed in 4% paraformaldehyde at room temperature for 30 minutes. The cultures were then washed 3 times and the transwell supports were excised for staining. The cells were permeabilized with 0.2% Triton X-100 in PBS for 10 minutes and then blocked with 10% normal donkey serum and 1% BSA for 60 minutes at room temperature. Primary antibody incubation was done overnight at 4 ºC followed by secondary incubation with Alexa Fluor® dyes for 60 minutes at room temperature. See **Table S1** for the manufacturer and dilution used for each antibody. Confocal images were acquired using the Olympus Fluoview System (Z-axis step 0.5μm; sequential scanning).

### Trans-epithelial electrical resistance (TEER) measurement

TEER was quantified using an EVOM ohm-voltmeter (World Precision Instruments, Sarasota, Fl) as previously described. In brief: transwells were placed into the Endohm-6 measurement chamber with PBS supplemented with calcium and magnesium (cmPBS) in the basal compartment and 200 μL of cmPBS in the apical compartment. TEER measurements were converted to Ohms-cm^2 based on the surface area of the transwell inserts.

### Lactate dehydrogenase assay

Apical fluid samples collected in PBS were assayed for cytotoxicity via lactate dehydrogenase (LDH) assay. LDH release was quantified using an LDH Cytotoxicity Detection Kit (Roche) according to the manufacturer’s instructions. Apical fluid collected from mock-infected nasal cultures was used to subtract background signal. Percentage cytotoxicity was calculated relative to ceiling LDH release values (quantified from cultures treated with Triton-X 100).

### Quantitative PCR (qRT-PCR)

Cells were lysed at indicated time points with buffer RLT Plus (Qiagen RNeasy Plus) and RNA was extracted following the manufacturer’s protocol. RNA was reverse transcribed into complementary DNA (cDNA) using the High Capacity Reverse Transcriptase Kit (Applied Biosystems). This cDNA was amplified using specific qRT-PCR primers for each target gene, iQ SYBR Green Supermix (Bio-Rad), and the QuantStudio 3 PCR system (Thermo Fisher). Primer sequences were as follows: 18S (Fwd: TTCGATGGTAGTCGCTGTGC, Rev: CTGCTGCCTTCCTTGAATGTGGTA); ACE2 (Fwd: AGAACCCTGGACCCTAGCAT, Rev: AGTCGGTACTCCATCCCACA); DPP4 (Fwd: GAAAGGTGTCAGTACTATTCTGTGT, Rev: CCAGGACTCTCAGCCCTTTATC). ∆Ct values were calculated using the formula ∆Ct = Ct_gene of interest_ – Ct_18S_. ∆(∆Ct) were calculated by subtracting sham-treated ∆Ct values from ∆Ct values for IL-13-treated cultures. Technical triplicates were averaged, and changes in mRNA levels were reported as fold changes over sham-treated cultures, using the formula 2^−∆(∆Ct)^.

### Western blotting

Cell lysates were harvested at indicated time points with RIPA buffer (50mM Tris pH 8, 150mM NaCl, 0.5% deoxycholate, 0.1% SDS, 1% NP40) supplemented with protease inhibitors (Roche: cOmplete mini EDTA-free protease inhibitor) and phosphatase inhibitors (Roche: PhosStop easy pack). Lysates were harvested via scraping of the transwell insert and incubated on ice for 20 minutes, centrifuged for 10 min at 15,000 RPM at 4ºC, and the supernatant was mixed 3:1 with 4X Laemmli sample buffer. Samples were boiled at 95ºC for 5 minutes, then separated via SDS/PAGE and transferred to polyvinylidene difluoride membranes. Blots were blocked in either 5% nonfat milk or 5% BSA in TBST and probed with antibodies as listed in **Table S2**. Blots were visualized using Thermo Scientific SuperSignal West Femto Substrate (catalog # 34096). Blots were stripped using Thermo Scientific Restore Western Blot stripping buffer (catalog #21059) for one hour at room temperature and then probed sequentially with antibodies.

## ACKNOWLEDGEMENTS

We thank members of the Weiss lab for feedback and discussion of this project, Dr. Nikki Tannetti for reading the manuscript, and Dr. Anthony Fehr for construction of the MERS-nsp15/ΔNS4a mutant. We also thank Drs. David W. Kennedy, James N. Palmer, Nithin D. Adappa, and Michael A. Kohanski for aid in the collection of nasal tissue for establishing primary nasal epithelial cultures. This work was supported by National Institutes of Health grants R01 AI140442 (SRW) and R01AI169537 (SRW&NAC); Department of Veterans Affairs Merit Review 1-I01-BX005432-01 (NAC&SRW); the Penn Center for Research on Coronaviruses and Other Emerging Pathogens (SRW). CO was supported in part by T32 AI055400 and AF in part by T32 AI007324.

