## Supplementary material for "Infection of primary nasal epithelial cells differentiates among lethal and seasonal human coronaviruses": supplemntal tables and figures

**Supplementary Materials**

Table S1
Table S2

Legends for supplementary figures 1-3
Figure S1
Figure S2
Figure S3

**Table S1. Antibodies used for western blotting**

| Primary Antibody | Antibody species | Blocking buffer | Dilution | Catalog number |
|---|---|---|---|---|
| DPP4/CD26 | Rabbit | 5% BSA/TBST | 1:1000 | Cell Signaling Technology 67138S |
| ACE2 | Rabbit | 5% milk/TBST | 1:1000 | Cell Signaling Technology 4355S |
| MUC5AC | Mouse | 5% milk/TBST | 1:500 | Sigma-Aldrich AMAB91539 |
| Type IV β-tubulin | Rabbit | 5% milk/TBST | 1:1000 | Abcam ab179509 |
| SARS-CoV-2 Nucleocapsid | Rabbit | 5% milk/TBST | 1:2000 | Genetex GTX135357 |
| MERS-CoV Nucleocapsid | Mouse | 5% milk/TBST | 1:2000 | Sino Biological 40068-MM10 |
| HCoV-NL63 Nucleocapsid | Rabbit | 5% milk/TBST | 1:2000 | Sino Biological 40641-T62 |
| GAPDH | Rabbit | 5% milk/TBST | 1:2000 | Cell Signaling Technology 2118S |
| **Secondary Antibody** | **Antibody species** | **Blocking buffer** | **Dilution** | **Catalog number** |
| Anti-rabbit IgG HRP-linked | Goat | Same as primary | 1:3000 | Cell Signaling Technology 7074S |
| Anti-mouse IgG HRP-linked | Goat | Same as primary | 1:3000 | Cell Signaling Technology 7076S |

**Table S2. Antibodies used for immunofluorescence assay**

| **Primary Antibody** | **Antibody species** | **Dilution** | **Catalog number** |
|---|---|---|---|
| DPP4/CD26 | Rabbit | 1:1000 | Cell Signaling Technology 67138S |
| MUC5AC | Mouse | 1:1000 | Sigma-Aldrich AMAB91539 |
| Type IV β-tubulin | Rabbit | 1:1000 | Abcam ab179509 |
| SARS-CoV-2 Nucleocapsid | Rabbit | 1:1000 | Genetex GTX135357 |
| MERS-CoV Nucleocapsid | Mouse | 1:1000 | Sino Biological 40068-MM10 |
| HCoV-NL63 Nucleocapsid | Rabbit | 1:1000 | Sino Biological 40641-T62 |
| **Secondary Antibody** | **Species** | **Dilution** | **Catalog number** |
| Alexa Fluor secondary dyes | Donkey | 1:1000 | Thermo Fisher |

**Figure S1 Immunofluorescence imaging controls in uninfected cultures**. (A) Uninfected nasal ALI cultures were stained with ciliated cell marker type IV β-tubulin (green), goblet cell marker mucin MUC5AC (red), and Hoescht (blue) to illustrate typical morphology prior to infection. (B) Uninfected cultures were stained with primary antibodies against the nucleocapsid protein for each of SARS-CoV-2, MERS-CoV, and HCoV-NL63.

**Figure S2 Trans-epithelial electrical resistance for mock-infected ALI cultures over time.** Duplicate nasal ALI cultures derived from each of 4 donors were mock-infected and TEER quantified at baseline, as well as at 96 and 192 hours post mock-infection. TEER values for each individual transwell are connected at each time point in order to monitor changes in TEER over time.

**Figure S3 HCoV replication is significantly impacted by IL-13 treatment of nasal epithelial cultures.** Nasal ALI cultures derived from 10 donors (for SARS-CoV-2 and MERS-CoV) or 7 donors (for HCoV-NL63) were sham- or IL-13-treated for the final 2 weeks of differentiation and then were infected in triplicate with each HCoV. ASL was collected at 48 and 96 hpi and infectious virus quantified via plaque assay. (A-C) Average viral titer at 48 hpi for sham- and IL-13-treated cultures for each donor were plotted as box-and-whisker plots, depicting the overall mean and interquartile range in titers for each condition. (D-E) Similar box-and-whisker plots comparing titers at 96 hpi in donor-matched sham- vs. IL-13 treated cultures. Note that these data are equivalent to data depicted in Figure 8, plotted differently. Statistical significance of difference in titer in sham vs. IL-13 treated cultures was determined for each HCoV at each time point using paired t-tests: **, $P \leq 0.01$; ***, $P \leq 0.001$; ****, $P \leq 0.0001$.

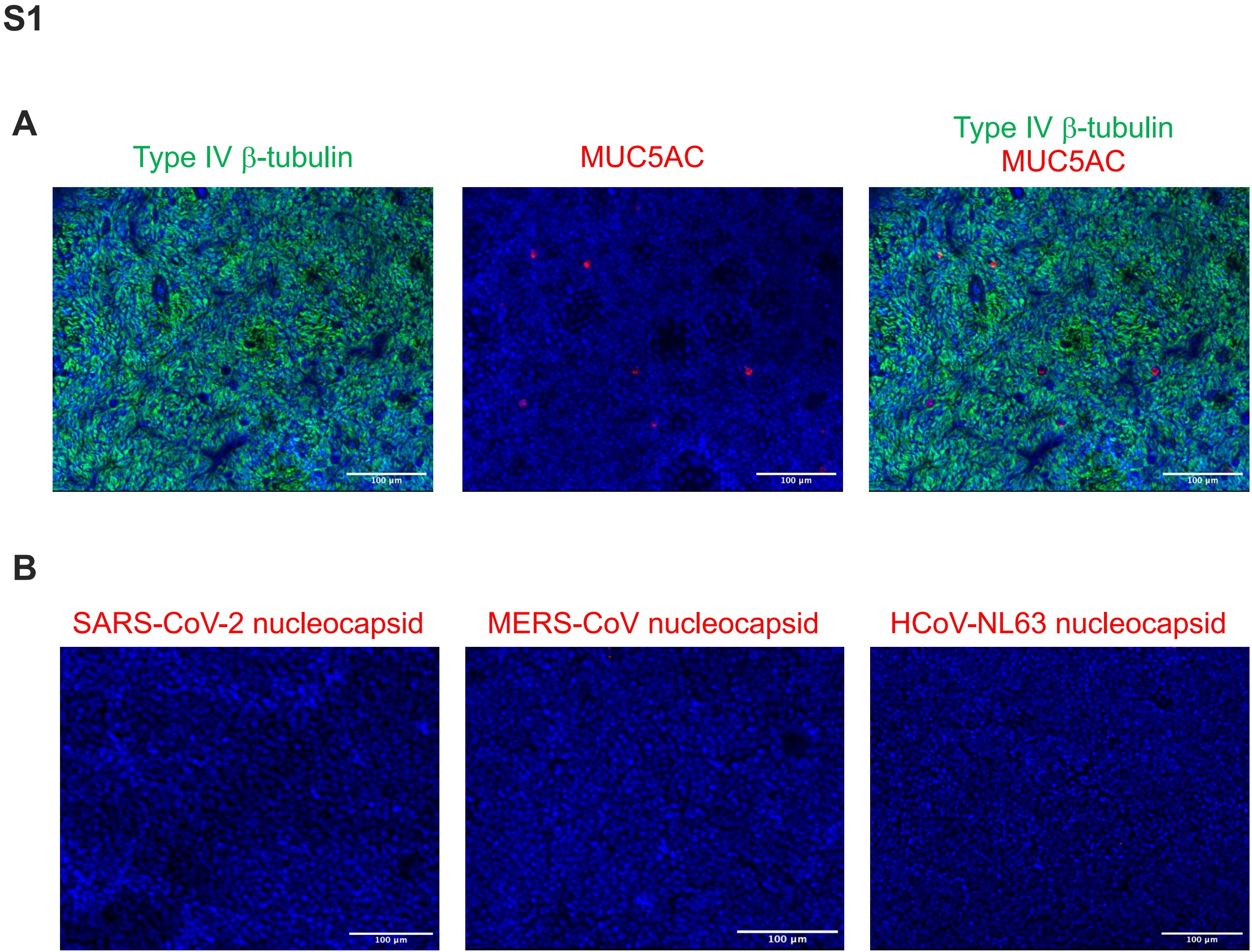
S1
A
Type IV β-tubulin
MUC5AC
Type IV β-tubulin
MUC5AC
100 µm
100 µm
100 µm
B
SARS-CoV-2 nucleocapsid
MERS-CoV nucleocapsid
HCoV-NL63 nucleocapsid
100 µm
100 µm
100 µm

**S2**

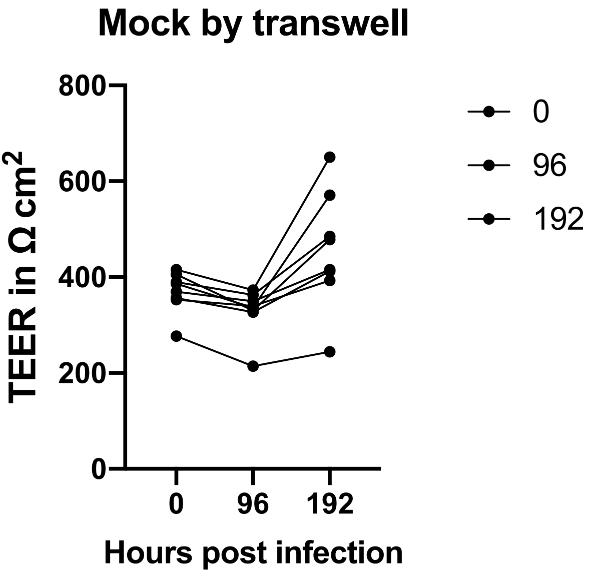

Mock by transwell
TEER in Ω cm2
800
600
400
200
0
0
96
192
Hours post infection
0
96
192

S3

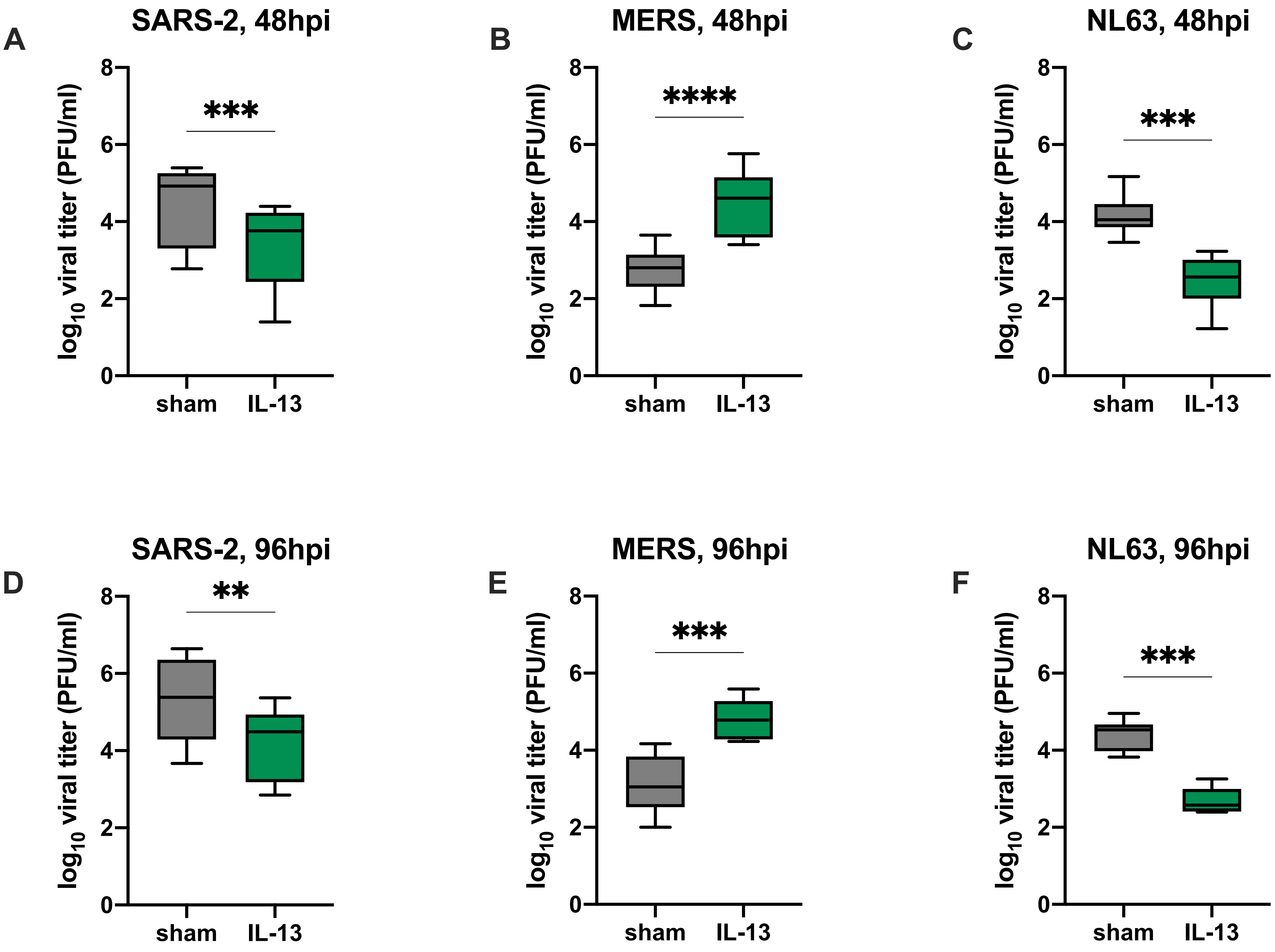
A
SARS-2, 48hpi
***
B
MERS, 48hpi
****
C
NL63, 48hpi
***
D
SARS-2, 96hpi
**
E
MERS, 96hpi
***
F
NL63, 96hpi
***
log10 viral titer (PFU/ml)
0
2
4
6
8
sham
IL-13
